# A 30-minute nucleic acid amplification point-of-care test for genital *Chlamydia trachomatis* infection in women: a prospective, multi-centre study of diagnostic accuracy

**DOI:** 10.1101/196675

**Authors:** EM Harding-Esch, EC Cousins, S-LC Chow, LT Phillips, CL Hall, N Cooper, SS Fuller, AV Nori, R Patel, S Thomas-William, G Whitlock, SJE Edwards, M Green, J Clarkson, B Arlett, JK Dunbar, CM Lowndes, ST Sadiq

## Abstract

**Background:** Rapid Point-Of-Care Tests (POCTs) for *Chlamydia trachomatis* (CT) may reduce onward transmission and reproductive sexual health (RSH) sequelae by reducing turnaround times between diagnosis and treatment. The io^^®^^ single module system (Atlas Genetics Ltd) runs clinical samples through a microfluidic CT cartridge, delivering results in 30 minutes. We evaluated its performance on female genital samples in four UK Genito-Urinary Medicine (GUM)/RSH clinics.

**Methods:** Prospective diagnostic accuracy study, using BD ProbeTec CT/GC assay as the routine clinic nucleic acid amplification test (NAAT) as the initial comparator test, and the QIAgen Artus CT assay to resolve discrepancies. In these instances, the reference standard was defined as the resolved result when two out of three assay results concurred. Female participants aged ≥16 provided additional-to-routine self-collected vulvovaginal swabs. Samples were tested fresh with the io^^®^^ CT assay within 7 days of collection, or were frozen at −80°C for later testing. Participant clinical, demographic and behavioural characteristics were collected to assess risk factors associated with CT infection.

**Results:** Of 785 participants recruited, final analyses were conducted on 709 (90.3%). CT prevalence was 7.2% (51/709) overall. Sensitivity, specificity, positive and negative predictive values of the io^^®^^ CT assay were, respectively, 96.1% (95% Confidence Interval (CI): 86.5-99.5), 97.7% (95%CI: 96.3-98.7), 76.6% (95%CI: 64.3-86.2) and 99.7% (95%CI: 98.9-100). There was no significant difference in performance measures between fresh and frozen samples, or between symptomatic and asymptomatic participants (p>0.05). The only risk factor associated with CT infection was being a sexual contact of an individual with CT.

**Conclusions:** The io^^®^^ CT-assay is the only 30-minute, fully automated, high-performing NAAT currently CE-marked for CT diagnosis in women, making it a highly promising diagnostic to enable specific treatment, initiation of partner notification and appropriately intensive health promotion at the point of care. Future research is required to evaluate acceptability by clinicians and patients in GUM/RSH clinics, impact on clinical pathways and patient management, and cost-effectiveness.

## BACKGROUND

Genital infection with *Chlamydia trachomatis* (CT) is a major public health challenge with over 200,000 CT diagnoses made in England alone in 2016, accounting for nearly half of all new sexually transmitted infection (STI) diagnoses that year ^1^. CT infection, which most commonly occurs in young people aged 15-24 years, goes undiagnosed in a large proportion of cases, is often asymptomatic in women (70%) and men (50%), and can lead to serious reproductive health morbidities, such as pelvic inflammatory disease, tubal infertility and ectopic pregnancy ^2^. The treatment of new CT diagnoses alone was predicted to contribute to nearly a third of the estimated direct medical costs of treating new STI diagnoses in the UK in 2011 ^3^.

Shortening the duration of infectiousness (between becoming infected and receiving effective treatment) of CT in at-risk individuals, is key to curbing CT transmission ^4-5^, while shortening the duration of infection can reduce reproductive health complications ^6-7^. The time between testing and treatment can vary widely ^8^ meaning some patients will wait longer for results and therefore have increased risk of transmitting infection. Public health programmes, such as the National Chlamydia Screening Programme (NCSP) in the UK, set standards for the time to treatment to guide services and reduce variation in care, thereby bringing the average time to treatment down. Nucleic acid amplification tests (NAATs) are recommended for routine diagnosis of CT infections by national and international guidelines ^9-11^ because of their high sensitivity and specificity, ability to deliver high volume testing and their relatively low cost. In addition to these advantages, NAAT-based point-of-care tests (POCTs), which enable patients to be tested and treated within the same clinical visit, have the potential to reduce the time to treatment and improve patient care ^12–14^. Rapid NAATs, such as the Cepheid CT/*Neisseria gonorrhoeae* (NG) GeneXpert assay, have equivalent performance characteristics to traditional lab-based NAATs^15^, but the 90 minute test-run time is too long for the test to be considered a POCT in many healthcare settings^14-16-17^. In addition, studies have shown a significant proportion of service users are unwilling to wait more than around 20 minutes for test results ^18-19^. There is therefore a need for more rapid (≤30 minutes) NAAT-based POCTs to aid accurate and specific diagnoses during one clinic visit.

The Atlas Genetics io^^®^^ platform is a 30-minute NAAT, single module system. Clinical samples, such as swab eluate, are transferred directly into a microfluidic cartridge for the diagnosis of genital CT in females. The objective of this study was to conduct a performance evaluation of the io^^®^^ CT assay using genital samples collected from females attending UK Genito-Urinary Medicine (GUM)/Reproductive and Sexual Health (RSH) clinics. Secondary objectives were to assess factors associated with being CT positive to inform on how to potentially implement the io^^®^^ CT assay in clinical pathways.

## METHODS

Four GUM/RSH clinics in London (Dean Street Clinic, Chelsea and Westminster Hospital NHS Trust), Taunton (Musgrove Park Hospital, Taunton and Somerset NHS Foundation Trust), Portsmouth (Solent NHS Trust) and Stevenage (Kingsway Sexual Health Service, Central London Community Healthcare NHS Trust) were selected for participant recruitment, which took place between June 2015 and March 2016. This manuscript was written following STARD (Standards for the Reporting of Diagnostic accuracy) guidelines ^20^ (supplementary material; files 1 and 2).

### Participants

Symptomatic and asymptomatic female participants were recruited prospectively and were considered eligible for the evaluation if they were: attending the participating GUM/RSH clinics and having a routine CT NAAT; aged 16 years or over; able to provide written informed consent; able and willing to provide an additional to routine self-collected vulvovaginal swab (SCVS). We defined participants to be potentially symptomatic for CT if they presented with any of the following: genital itching, discharge (clear or cloudy liquid from the vagina), pain/burning when urinating, needing to pass urine more often than usual, pain deep inside the vagina when having sex, pain just inside or around the vagina when having sex, bleeding after sex, bleeding in between periods or pelvic abdominal pain. Participants were recruited following provision of written informed consent and given a unique participant identifier (participant ID). Routine clinical and demographic data were recorded prospectively by a delegated clinical staff member.

Based on a conservative estimate of the sensitivity of the io^^®^^ CT assay of 92% (95% confidence interval (95%CI); 81-97) and a mean CT positive rate of 7.5% in GUM/RSH clinics (Hamish Mohammed, Public Health England (PHE), personal communication) the target sample size was 750 females to obtain 50 CT positive female samples. Recruitment of participants formed a convenience sample, with clinics preferentially recruiting participants who they judged were more likely to be CT positive (e.g. CT-positive sexual partner) ^21^ in order to increase the likelihood of meeting the 50 CT positive target.

### Specimen collection

Following collection of the clinic’s routine vulvovaginal swab for CT/NG NAAT diagnosis, an SCVS (Copan eNAT^^®^^ Collection and Preservation System) was provided for the study. If the participant was having a vaginal examination, she was asked to take the SCVS sample before the examination. Routine vulvovaginal swabs were processed as per clinical protocol, and the additional SCVS samples stored at room temperature, initially in clinic for a maximum of six days, and then transferred at between 2-8°C to the Applied Diagnostic Research and Evaluation Unit (ADREU) laboratories at St George’s, University of London (SGUL). Upon receipt, samples were aliquoted as follows: 600 μl for testing with the io^^®^^ CT assay; 300 μl for discrepant result testing, if necessary (see below); and any remainder (approximately 1ml) for repeat testing as required. Samples received with insufficient volume for both an initial and discrepant sample aliquot were excluded from the study. Samples were either tested “fresh” within seven days of collection (stored refrigerated (2-8°C) at SGUL) or immediately frozen at −80°C.Frozen samples were defrosted for a minimum of 30 minutes and tested within two hours.

The research sample and case report form data were linked to clinical results using the unique participant IDs. Once all data were matched and data verification complete, the temporarily list linking participant and clinical identifiers was destroyed, thus anonymising the data.

### Test methods

Four io^^®^^ systems were used to test the samples between September 2015 and September 2016. io^^®^^ CT assay cartridges were kept refrigerated prior to use. Positive and negative io^^®^^ CT assay control cartridges were run on each io^^®^^ system daily before sample testing to validate the system. A fixed volume pipette, packaged with each cartridge, was used to withdraw and transfer 500 μl of sample into the cartridge, which was then loaded onto the io^^®^^ system. The participant ID was scanned into the system and testing started via a touch-screen control. In all cases, results were delivered in 30 minutes as either ‘CT detected’, ‘CT not detected’ or ‘invalid’. If a sample returned an invalid result, a repeat test was performed on a new cartridge. If invalid a second time, the final result was recorded as invalid. ADREU laboratory staff carrying out the testing on the io^^®^^ system were blind to participant clinical information and the clinic CT/NG NAAT results.

### Estimating diagnostic accuracy

For all samples, the initial comparator test used was the CE-marked Becton Dickinson (BD) ProbeTec^™^ Qx CT/GC assay (Oxford, UK), run on the BD Viper analyser, as this was the routine CT/NG NAAT used at all participating GUM/RSH clinics. Those conducting the initial comparator test were blind to clinical information and io^^®^^ CT assay results. The io^^®^^ CT assay results were compared to the initial comparator test results by the ADREU study Coordinator, and any discordant results identified.

We defined the reference standard ^20^ as the initial comparator test result when in agreement with the io^^®^^ CT assay result. If the io^^®^^ CT assay result did not agree with the initial comparator test result, a further test with the CE-marked QIAgen Artus^^®^^ *C. trachomatis* Plus RG PCR kit (Manchester, UK), run on the Qiagen Rotor-Gene Q 2plex HRM PCR thermocycler, was performed according to manufacturer’s instructions. In these cases the reference standard was defined as the resolved result when two out of three of io^^®^^ CT assay, initial reference test and Artus CT assay results were in agreement. This discrepant analysis approach was employed as a result of budgetary and time constraints.

### Statistical analysis

Data from the io^^®^^ system were transcribed manually onto the study database. Data cleaning and validation were performed independently and separately at SGUL and at PHE. Any discrepancies were resolved through checking the original data with the clinics. Data were analysed at PHE using Stata (StataCorp LP v13.1). Missing data were verified, and all initial comparator test results double-checked, with each clinic. Participants with either a missing io^^®^^ CT assay or initial comparator test result were excluded from analyses.

Diagnostic accuracy metrics (sensitivity, specificity, positive (PPV) and negative (NPV) predictive values) and their 95% CIs (binomial exact) were calculated. A two-sample chi-squared test was performed to compare results by symptomatic and asymptomatic status and sample storage method (frozen versus unfrozen (“fresh”)) (Table 1). CT prevalence was assessed at the clinic population level using the GUM Clinical Activity Dataset (GUMCAD) ^22^ for the clinics involved over the study period to inform if there had been any bias in recruitment. We also compared the impact of the performance measures of the io^^®^^ CT assay on PPV and NPV using national GUMCAD prevalence data (Hamish Mohammed, PHE, personal communication) (high prevalence setting) as well as from Natsal-3 (National Survey of Sexual Attitudes and Lifestyles) ^23^ (low prevalence setting), focusing on women aged 16-24 years. Natsal-3 is a population-based prevalence survey representing all adults resident in the UK between the ages of 16-74 years.

**Table 1.** Diagnostic accuracy of the io^^®^^ CT assay when compared with the reference standard^a^.

|  |  |  |  |  |  |  |  |
| --- | --- | --- | --- | --- | --- | --- | --- |
|  | All | Symptomatic | Asymptomatic | Symptomatic vs asymptomatic p-value<sup>b</sup> | Fresh | Frozen | Fresh vs frozen p-value<sup>b</sup> |
| Prevalence | 7.2<br>(5.4-9.3)<br>(51/709) | 11<br>(6.0-18.1)<br>(13/118) | 6.4<br>(4.6-8.7)<br>(38/591) | 0.106 | 9.3<br>(6.1-13.3)<br>(26/281) | 5.8<br>(3.8-8.5)<br>(25/428) | 0.111 |
| Sensitivity | 96.1<br>(86.5-99.5)<br>(49/51) | 100.0<br>(75.3-100.0)<br>(13/13) | 94.7<br>(82.3-99.4)<br>(36/38) | 0.906 | 96.2<br>(80.4-99.9)<br>(25/26) | 96.0<br>(79.6-99.9)<br>(24/25) | 0.997 |
| Specificity | 97.7<br>(96.3-98.7)<br>(643/658) | 98.1<br>(93.3-99.8)<br>(103/105) | 97.6<br>(96.0-98.7)<br>(540/553) | 0.976 | 96.5<br>(93.4-98.4)<br>(246/255) | 98.5<br>(96.8-99.5)<br>(397/403) | 0.854 |
| PPV | 76.6<br>(64.3-86.2)<br>(49/64) | 86.7<br>(59.5-98.3)<br>(13/15) | 73.5<br>(58.9-85.1)<br>(36/49) | 0.706 | 73.5<br>(55.6-87.1)<br>(25/34) | 80.0<br>(61.4-92.3)<br>(24/30) | 0.824 |
| NPV | 99.7%<br>(98.9-100.0)<br>(643/645) | 100.0<br>(96.5-100.0)<br>(103/103) | 99.6<br>(98.7-100.0)<br>(540/542) | 0.981 | 99.6<br>(97.8-100.0)<br>(246/247) | 99.7<br>(98.6-100.0)<br>(397/398) | 0.989 |

CT, *Chlamydia trachomatis*; PPV, Positive Predictive Value; NPV, Negative Predictive Value
<sup>a</sup> Values are percentages (95% confidence intervals) (numbers)
<sup>b</sup> Study prevalence of CT and performance characteristics (sensitivity, specificity, PPV and NPV) were compared between symptomatic and asymptomatic participants and between fresh and frozen samples using the Pearson Chi-squared test for comparing two proportions.

Univariate logistic regression analysis of risk factors associated with CT infection (as defined by the reference standard) was conducted. Factors considered significant (p<0.05) were included in a multivariate analysis employing a forward step-wise approach, with age and clinic considered *a priori* risk factors. The same process was used to determine any factors associated with an invalid io^^®^^ CT assay result. Factors included in the analyses were: participant age, sexual orientation, having taken medication that would be active against CT infection in the last 6 weeks, being a contact of a CT-positive individual, having had an STI in the last 12 months, whether currently menstruating, whether symptomatic for CT infection, and clinic attended. In the invalid io^^®^^ CT assay result logistic regression analysis, clinic routine CT NAAT and NG NAAT results were also included as explanatory variables.

## RESULTS

### Participants and sites

A total of 785 female participants were recruited from the different clinics (Figure 1). 76 participants were excluded from the final analyses, conducted on 709 (90.3%) participants, for reasons including not fulfilling eligibility criteria, missing initial reference test data (clinic BD Viper CT/NG NAAT), and final invalid io^^®^^ CT assay results. Overall CT prevalence according to the reference standard among the samples tested was 7.2% (51/709; 95%CI 5.4-9.3) with no difference between fresh and frozen samples (Table 1). Baseline participant demographic and clinical characteristics are presented in Table 2.

**Table 2.** Risk factor analysis for being CT positive^a^.

| Table 2. Risk factor analysis for being CT positive<sup>a</sup> |  |  |  |  |  |  |  |  |
| --- | --- | --- | --- | --- | --- | --- | --- | --- |
|  |  |  | Univariate analysis |  |  | Multivariate analysis<sup>c</sup> |  |  |
| Characteristic | No. of participants | No. (%) with CT<sup>b</sup> | OR | 95%CI | P-value | OR | 95%CI | P-value |
| Age | 709 | 51 (7.2) | 0.94 | 0.89 - 0.99 | 0.01 | 0.95 | 0.90 - 1.00 | 0.06 |
| Clinic |  |  |  |  |  |  |  |  |
| 1 | 387 | 34 (8.8) | 1 |  |  | 1 |  |  |
| 2 | 92 | 3 (3.3) | 0.35 | 0.11 - 1.17 | 0.09 | 0.36 | 0.10 - 1.29 | 0.12 |
| 3 | 72 | 2 (2.8) | 0.30 | 0.07 - 1.26 | 0.10 | 0.31 | 0.07 - 1.43 | 0.13 |
| 4 | 158 | 12 (7.6) | 0.85 | 0.43 - 1.69 | 0.65 | 0.47 | 0.20 - 1.07 | 0.07 |
| Contact of a CT positive |  |  |  |  |  |  |  |  |
| No | 617 | 26 (4.2) | 1 |  |  | 1 |  |  |
| Yes | 48 | 21 (43.8) | 17.68 | 8.85 - 35.33 | <0.001 | 17.33 | 8.28 - 36.27 | <0.001 |
| Not known | 44 | 4 (9.1) | 2.27 | 0.76 - 6.83 | 0.14 | 2.94 | 0.91 - 9.45 | 0.07 |
| Taken CT-active medication in last 6 weeks |  |  |  |  |  |  |  |  |
| No | 669 | 47 (7.0) | 1 |  |  |  |  |  |
| Yes | 40 | 4 (10.0) | 1.47 | 0.50 - 4.31 | 0.48 |  |  |  |
| Symptomatic |  |  |  |  |  |  |  |  |
| No | 591 | 38 (6.4) | 1 |  |  |  |  |  |
| Yes | 118 | 13 (11.0) | 1.8 | 0.93 - 3.50 | 0.08 |  |  |  |
| STI last year |  |  |  |  |  |  |  |  |
| No | 641 | 41 (6.4) | 1 |  |  | 1 |  |  |
| Yes | 68 | 10 (14.7) | 2.52 | 1.20 - 5.30 | 0.01 | 2.09 | 0.87 - 4.99 | 0.10 |
| Currently menstruating |  |  |  |  |  |  |  |  |
| No | 634 | 46 (7.3) | 1 |  |  |  |  |  |
| Yes | 71 | 4 (5.6) | 0.76 | 0.27 - 2.19 | 0.62 |  |  |  |
| Has sex with<sup>d</sup> |  |  |  |  |  |  |  |  |
| Men | 665 | 47 (7.1) | 1 |  |  |  |  |  |
| Women | 20 | 1 (5.0) | 0.70 | 0.09 - 5.28 | 0.72 |  |  |  |
| Both | 23 | 3 (13.0) | 1.98 | 0.57 - 6.88 | 0.29 |  |  |  |

CT, *Chlamydia trachomatis*; OR, Odds ratio; 95% CI, 95% Confidence Interval; STI, Sexually transmitted infection
<sup>a</sup> For each characteristic the number of participants and the proportion of these with a CT infection are shown.
<sup>b</sup> CT positive defined as reference standard: either positive initial comparator test result (when in agreement with io<sup>®</sup> CT assay result); or positive by at least two of three of the initial comparator test, io<sup>®</sup> CT assay and, Artus CT assay).
<sup>c</sup> Adjusted for age, clinic, contact status and STI in the last year (age and clinic considered *a priori* risk factors and included in all models).
<sup>d</sup> Sexual orientation unknown for one participant.

**Figure 1.**
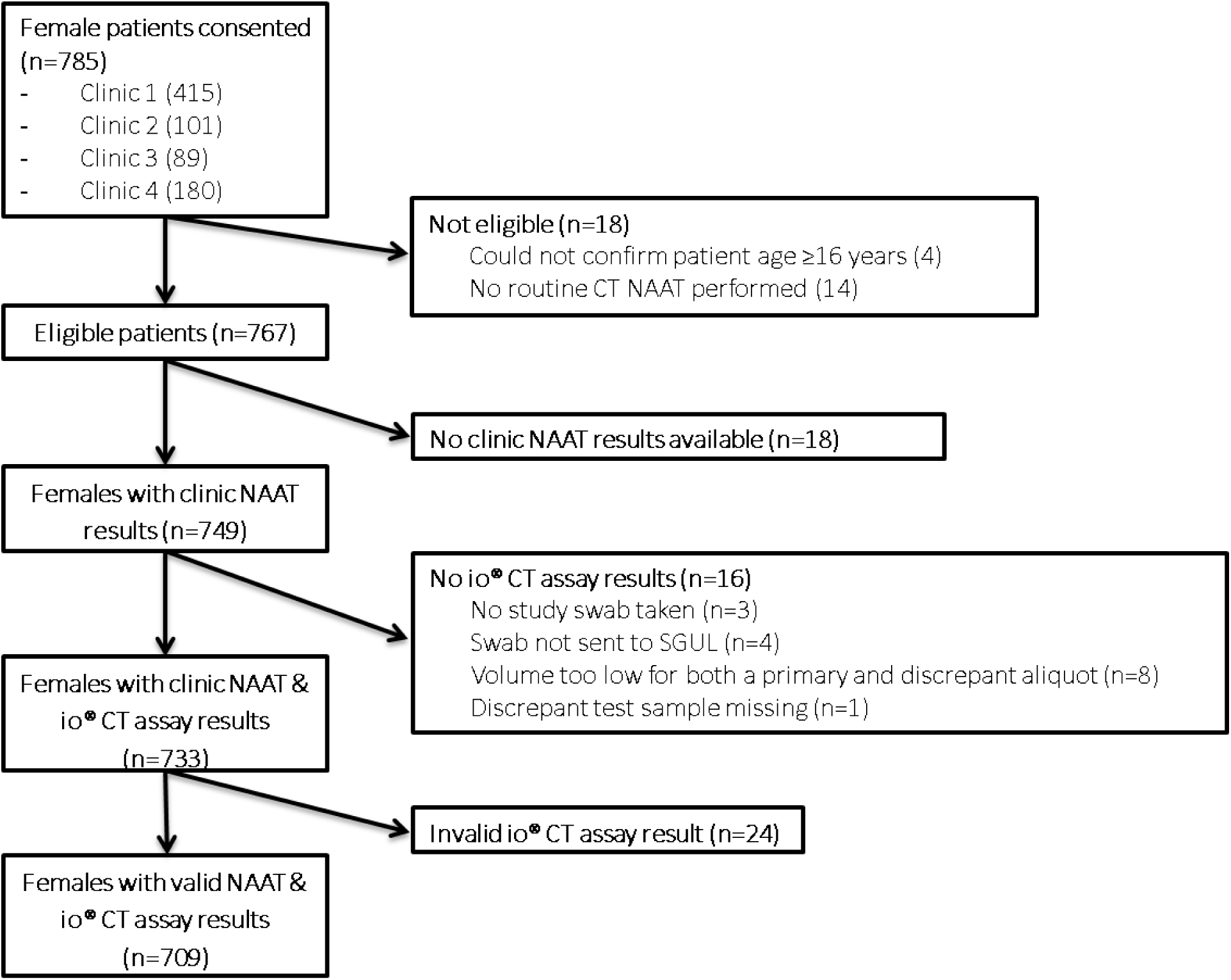
Flow chart summarising patient recruitment and sample collection results availability. CT, *Chlamydia trachomatis*; NAAT, Nucleic Acid Amplification Test Flow diagram showing total number of eligible participants who consented to the study, ending with the total number of participants included in the final analyses. Samples were excluded where the participant did not meet the study eligibility criteria (n=18); that did not have a clinic NAAT result available (n=18), that were not tested on the io^^®^^ CT assay (n=16); or that had a final invalid result by the io^^®^^ CT assay (n=24).

### Performance of the io^^®^^ CT assay

A number of tests (n=24/733; 3.3%) reported an invalid result on the first run and 100% of these reported an invalid result on a second. Of the 24 invalid results 20 were negative and four were positive for CT by the clinic routine CT/NG NAAT. In the logistic regression analysis, there were no factors associated with the io^^®^^ CT assay result being invalid. 684/709 (96.5%) io^^®^^ CT assay results agreed with the initial comparator test result. For the remaining 25 samples that had a discordant result between the io^^®^^ CT assay and initial comparator test, the Artus CT assay (used for discrepant testing) agreed with 8 io^^®^^ CT assay results (4/19 io^^®^^ CT assay positives and 4/6 io^^®^^ CT assay negatives). Resulting sensitivity, specificity, PPV and NPV overall, compared with the reference standard, were 96.1% (95%CI 86.5-99.5), 97.7% (95%CI 96.3-98.7), 76.6% (95%CI 64.3-86.2) and 99.7% (95%CI 98.9-100.0) respectively (Table 1). There were no significant differences in the performance of the io^^®^^ CT assay in any of the diagnostic accuracy measures between symptomatic and asymptomatic participants, or between fresh and frozen SCVS samples run on the io^^®^^ system (Table 1).

The study CT prevalence of 7.2% was higher than the 5.2% (p=0.0182) prevalence of all female patients routinely tested for CT attending the study clinics over the study period, determined by clinic GUMCAD data ^22^. National CT prevalence data for females from GUMCAD (Hamish Mohammed, PHE, personal communication), and Natsal-3 (Sept 2010-Aug 2012) ^23^ were 6.7% and 3.1%, respectively. PPVs for these data sets were 74.4% (95%CI 65.8-81.5) and 57.7% (95%CI 47.3-67.4) respectively. NPVs were comparable to the study NPV.

### Risk factors for being CT positive

Factors associated with being CT positive in univariate logistic regression analysis were young age, being a sexual contact of a CT positive individual and having had an STI in the last year (Table 2). In multivariate analysis, only being a sexual contact of CT remained as an independent risk factor.

## DISCUSSION

In this first diagnostic performance evaluation of the io^^®^^ CT assay, a 30-minute NAAT POCT for detection of genital CT infection in symptomatic and asymptomatic women, we have shown sensitivity and specificity of 96.1% (95%CI 86.5-99.5) and 97.7% (95%CI 96.3-98.7) respectively. The PPV and NPV were 76.6% (95%CI 64.3-86.2) and 99.7% (95%CI 98.9-100.0) respectively, with a study CT prevalence of 7.2% (95%CI 5.4-9.3).

National and international guidelines recommend laboratories use NAATs for the diagnosis of CT infections due to their superior performance compared with other diagnostic technologies ^9–11^. Performance evaluations for US Food and Drug Administration (FDA) approval use a Patient Infection Status (PIS) or composite gold standard study design, with data for some of the most commonly used laboratory-based NAATs on self-collected vaginal swabs indicating sensitivities between 96.5%-98.4% and specificities between 95.6%-99.2% ^24–27^. The Cepheid GeneXpert CT/NG rapid NAAT, which is not a traditional laboratory-based NAAT, has a reported sensitivity and specificity of 98.7% (95%CI 93.1-100) and 99.4% (95%CI 98.9-99.7) using a PIS reference standard ^15^. We however employed a discrepant analysis approach for our study, and published evaluations of laboratory-based NAATs for CT detection using vaginal swabs using this approach report sensitivities of 80.4%-100.0%, and specificities of 99.5%-100.0% ^28–30^. The io^^®^^ CT POCT, as evaluated in our study, thus has a comparable sensitivity but potentially lower specificity than these laboratory-based NAATs. However, a previous evaluation of the io^^®^^ CT assay, also employing a discrepant analysis approach, indicated a specificity comparable to those reported for laboratory-based NAATs and higher than in our study, of 99.0% (USA) ^31^. This variation in performance measures achieved supports the importance of conducting diagnostic accuracy studies in different settings and populations ^32^.

A test’s PPV is directly affected by the CT prevalence in the population tested. To compare how the laboratory-based NAATs evaluated by discrepant analysis would perform in our study population, we calculated their PPVs using our study prevalence of 7.2%. The resulting PPVs ranged between 93.9% (95%CI 84.6-98.8) and 100% (95%CI 93.0-100.0). The io^^®^^ CT assay PPV in our study of 76.6% (95%CI 64.3-86.2) is at the lower end of the range. When applying the sensitivity and specificity estimates from the previous io^^®^^ CT assay diagnostic accuracy evaluation ^31^ to our study prevalence, a PPV of 87.9% (95%CI 75.5-94.7) was achieved. We also assessed how the io^^®^^ CT assay would perform in high (GUMCAD) and low (Natsal) CT prevalence settings by calculating how the predictive values would change based on the prevalence data reported for these settings. PPVs were 74.4% and 57.7% for GUMCAD and Natsal respectively, and NPVs were 99.7% and 99.9% respectively. Previous British Association for Sexual Health and HIV (BASHH) guidelines from 2010 ^33^ stated that testing platforms must have a PPV of over 90%. This is no longer mentioned in current BASHH guidelines ^34^, possibly because previous guidelines were written at a time when performance characteristics for laboratory-based NAATs were variable and when POCT NAATs were not available. The way in which the io^^®^^ CT assay is best implemented in clinical pathways as a POCT in view of these PPV results should be considered, for example assessing patient CT infection risk to increase CT positivity in the tested population.

Consequently, risk factor analyses may be helpful in targeting who to test with the io^^®^^ CT assay. Previous work in UK GUM/RSH clinics has indicated that younger age (<20), more than one (concurrent) sexual partner, black ethnicity and smoking are independent risk factors for CT in women ^35^. In our analysis risk factors included younger age, having had an STI in the last year and being a sexual contact of someone diagnosed with CT in univariate analyses, although only the latter remained an independent risk factor in the multivariate analysis. Being a sexual contact of an individual with a sexually transmitted infection is reported to be a risk factor for CT infection in studies from Europe and the US ^21^. It is important to consider that in other populations or settings these risk factors may be different ^21-23-36^, and targeted testing might need to be adjusted accordingly. This targeted-patient approach is further supported by the io^^®^^ system’s current single-modular platform design, which has the potential for multiple systems to be placed in a clinic at any one time.

Equally, there are practical implications of the io^^®^^ CT assay 3.3% failure rate, which although consistent with that of the GeneXpert CT/NG assay on first attempt ^15^, did not improve after repeat testing, suggesting that following an initial invalid result, patients would need to be recalled to provide a new sample. This may have implications for the cost-effectiveness of deploying this test, and with no factors predicting likelihood of an invalid result, it is not possible to factor in adjustments to sample collection pathways to mitigate potential invalid test results.

Sampling from different UK GUM/RSH clinics enabled us to both target high-risk patients to achieve the positivity required for the study and capture different populations that are more representative of the GUM/RSH clinic attendees for the whole of the UK than would have been possible with a single-site study. The operators performing the io^^®^^ CT assay testing were blind to the initial comparator test results, and *vice versa*, as well as to participant clinical and demographic characteristics, and data analysis was conducted independently at PHE. All participating clinics used the same routine clinic NAAT to provide a CT diagnosis for participants ensuring consistency across the study. However, only one sample type for one anatomical site was evaluated. Whereas vulvovaginal swabs are routinely used in GUM/RSH clinics for NAAT diagnosis of urogenital CT infection, endo-cervical and urine samples are also commonly used ^15-37^, and there is increasing evidence for the importance of extra-genital sampling in women ^38-39^. When analysing GUMCAD data for the study clinics during the study period, the CT prevalence of all women being tested for was 5.2%, lower than our participant 7.2% prevalence. Although this indicates a recruitment bias, probably because patients considered more likely to be CT-positive (e.g. CT-positive sexual partner ^21^) were approached to participate in order to meet our 50 CT-positive sample size, there is no reason to suspect that this would have affected our estimates of the io^^®^^ CT assay’s sensitivity and specificity.

We employed discrepant testing to define our reference standard if the io^^®^^ CT assay result did not agree with that of the initial comparator test, rather than a composite gold standard approach where at least two reference tests are used together (typically searching for a different target), and a clear definition of a positive and a negative is provided ^40-41^. Discrepant testing is known to introduce an initial bias towards the index test (io^^®^^ CT assay) ^42-43^. A better reference standard, decided by, for example, three independent reference tests against which the index test is compared, would have provided a more accurate estimate of performance of the io^^®^^ CT assay ^41^. However, this was not logistically possible within our study.

The io^^®^^ CT assay has been CE-marked and licensed for use in Europe in women only ^44^. Future research is required to evaluate its performance in different settings, as well as in male participants. Although previous modelling work has demonstrated that a CT-only POCT could have an epidemiological impact and be cost-saving at the local level ^45^, dual CT/NG NAAT testing is most commonly employed in GUM/RSH settings in the UK. Therefore, further work is needed to assess the feasibility, acceptability and cost-effectiveness of a CT-only POCT in GUM/RSH clinic settings. Atlas Genetics Ltd is currently developing a dual CT/NG assay for both male and female samples, which could overcome the limitations of a single-pathogen test.

## CONCLUSIONS

We have shown that the io^^®^^ CT-assay is the only 30-minute, fully automated, high-performing NAAT currently CE-marked for CT diagnosis in women, making it a highly promising diagnostic to enable specific treatment, initiation of partner notification and appropriately intensive health promotion at the point of care. Future research is required to evaluate the io^^®^^ CT assay’s acceptability by clinicians and patients in GUM/RSH clinics, impact on clinical pathways and patient management, and cost-effectiveness

## DECLARATIONS

### Ethics approval and consent to participate

This study was approved by the London Bridge Research Ethics Committee (REC reference 13/LO/0691, IRAS reference 126709). All participants gave written, informed consent before taking part in the study. Participants gave informed consent for anonymised results to be used by the researchers for publication in medical journals and presentation at scientific meetings.

### Funding

This study was funded by the UK Clinical Research Collaboration (Medical Research Council) Translation Infection Research Initiative Consortium (grant number G0901608) and by Atlas Genetics Ltd. This study was sponsored by St George’s, University of London. The sponsor had no role in the study design; the collection, analysis, or interpretation of data; the writing of the article; or the decision to submit it for publication.

The UKCRC had no role in the study design; in the collection, analysis, or interpretation of data; in the writing of the report; or in the decision to submit the article for publication. Atlas Genetics contributed to the conception of the study, reviewed the protocol for io^^®^^ platform methodology provided technical support during the study, and reviewed and commented on the final draft of the manuscript. Atlas Genetics did not have a role in data collection, analysis or interpretation.

### Authors’ contributions

EHE and EC contributed equally to this paper. EHE, STS, MG, and JC conceived the study. EHE, EC, SLCC, LTP, CLH, SSF, AVN, RP, STW, GW, SE, MG, JC, CML and STS designed and implemented the study. EHE, EC, LTP, NC, AVN, JKD and STS conducted data analyses. EHE, EC, NC and STS had full access to all of the data in the study, and take responsibility for the integrity of the data and the accuracy of the data analysis. All authors interpreted the data and critically revised the manuscript. EHE, EC and STS drafted the manuscript. STS is guarantor.

## Acknowledgements

The Applied Diagnostic Research and Evaluation Unit acknowledges the support of the National Institute for Health Research Clinical Research Network (NIHR CRN).

We are grateful to the participants and clinics for taking part.

Thanks to David Pearce, Dan Shenton and Stephanie Bannister from Atlas Genetics, for technical support.

Thanks to Hamish Mohammed from Public Health England, for providing GUMCAD data for sample size calculations.

